# South American *Plasmodium falciparum* isolates reveal new insights into the role of PfAAT1 in mediating resistance to Chloroquine

**DOI:** 10.64898/2026.07.15.738636

**Authors:** Kyle Michie, Ella Bishop, Vladimir Corredor, Diego F. Echeverry, Janet E. Deane, Julian C. Rayner

**Author notes:** These authors contributed equally to this work (Kyle Michie, Ella Bishop).

## Abstract

Antimalarial drug resistance remains one of the most significant challenges to global malaria control. The emergence of resistance to chloroquine, the first truly globally distributed antimalarial, has been extensively studied but is still not fully understood. While mutations in the *Plasmodium falciparum* chloroquine resistance transporter (PfCRT) primarily drive resistance, the phenotype is complex and multigenic, with mutations in the putative amino acid transporter PfAAT1 recently confirmed to play a modulatory role. To date studies have focused on PfAAT1 mutations found in African and Southeast Asian *P. falciparum* lineages, but chloroquine resistance emerged independently in South America, where there may be novel PfAAT1 polymorphisms that are functionally important. We used AlphaFold modelling to reveal high homology between PfAAT1 and the human lysosomal arginine transporter SLC38A9, which allows prediction of membrane orientation and identifies a partially open channel accessible from the cytoplasm. Several PfAAT1 mutations found only in South American isolates sit near the entrance of this pore, most notably V231 where mutation to aspartate is predicted to alter channel conformation and influence transport, while nearby P446A (which is always found in combination with V231D) and I248T are predicted to impact pore flexibility and local structural stability. To functionally validate these insights, we employed CRISPR/Cas9 gene editing across parasite strains with diverse geographic origins. Reverting the regional V231D mutation in the South American 7G8 strain significantly reduced CQ resistance, providing the first functional evidence that this residue modulates drug susceptibility. Furthermore, introducing the apparently Colombia-specific I248T mutation significantly enhanced parasite multiplication rates in 7G8, demonstrating complex fitness and sensitivity trade-offs. Our findings reinforce the distinct evolutionary trajectory for South American CQ resistance and highlight the necessity of including additional *pfaat1* mutations in global molecular surveillance strategies.

**Author Summary:** Antimalarial drug resistance is a major threat to global public health. Chloroquine was used widely in the 1950s-60s as part of a global malaria eradication campaign, but resistance emerged in multiple places independently and chloroquine resistant parasites directly led to the death of millions of children. Chloroquine resistance is primarily driven by mutations in the transporter PfCRT which are thought to increase export of chloroquine from the digestive vacuole, where chloroquine acts to prevent the ability of the parasite to digest haemoglobin as a source of energy. However, it is becoming increasingly clear that additional vacuolar transporters can also modulate chloroquine resistance. In this study we focused on mutations in the putative amino acid transporter 1 (PfAAT1) which are specific to South America, where chloroquine resistance emerged independently from Southeast Asia. By integrating AlphaFold structural predictions with CRISPR/Cas9 gene editing across geographically diverse parasite backgrounds we provide the first functional evidence that the region-specific V231D mutation significantly diminishes chloroquine resistance. These findings emphasise that chloroquine resistance in South America followed a unique trajectory and that PfAAT1 is involved in complex fitness and sensitivity trade-offs. This work emphasises the benefits of integrating diverse experimental and modelling approaches with global molecular surveillance.

## Introduction

Malaria is a human disease caused by parasites of the *Plasmodium* genus, with most severe malaria caused by *Plasmodium falciparum*. While the majority of *P. falciparum*-associated mortality is concentrated in Sub-Saharan Africa, the parasite is globally distributed and is found in multiple countries in South America^1^, where it probably arrived through the transatlantic slave trade^2,3^. In 2024 there were close to 191 million cases of *P. falciparum* globally, of which 145 000 were in South America. 46 000 of these South American *P. falciparum* cases were registered in Colombia alone, accounting for 32% of cases in the continent.

Antimalarial chemotherapy remains an essential part of the fight to control *Plasmodium falciparum* malaria, but the drugs employed have changed significantly across the past century as a result of the evolution and spread of antimalarial drug resistance. Artemisinin Combination Therapies (ACTs) are the current front-line treatment for *P. falciparum*, but prior to the introduction of ACTs, several different classes of antimalarials were used across the 20^th^ century until they were lost to resistance. The antimalarial with perhaps the longest period of use prior to the introduction of ACTs was chloroquine. Chloroquine accumulates in the metabolically-active digestive vacuole^4^ where it binds intermediate haemoglobin metabolites, thereby preventing their polymerization into inert non-toxic crystals and leading to cytotoxic effects for the parasite^5^. Chloroquine was first introduced relatively soon after World War II, and was used globally and extensively as part of the WHO malaria eradication campaign of the 1950s^6^. By the 1960s however, clinical failure of chloroquine had begun to emerge. Resistance emerged *de novo* independently in South America^7^, the South Pacific^8–10^, and Southeast Asia^11^. Resistance then spread from Southeast Asia into Sub-Saharan Africa where it caused a major increase in mortality through the 1980s^6^.

Decades of studies to uncover the molecular basis of chloroquine resistance revealed a central role for multiple membrane-spanning transporter proteins that localise to the digestive vacuolar membrane. The primary mediator of chloroquine resistance is *Chloroquine Resistance Transporter* (PfCRT, Pf3D7_0709000)^12^; mutant PfCRT haplotypes are sufficient for chloroquine resistance^13^ and widely used as molecular markers^14^. Resistance-associated polymorphisms in PfCRT are thought to lead to a gain-of-function in increasing chloroquine efflux from the vacuole^13,15^. *Multidrug Resistance Transporter 1* (PfMDR1, Pf3D7_0523000) is also proposed to play a role, with polymorphisms and copy number variation thought to impact chloroquine import^16–18^.

More recently, attention has focussed on the contribution of a third transporter, *Amino Acid Transporter 1* (PfAAT1, Pf3D7_0629500)^19^. PfAAT1 is a putative peptide transporter^20,21^ and like PfCRT and PfMDR1 is a multi-pass membrane protein localised in the digestive vacuolar membrane^22^. A role for PfAAT1 in chloroquine resistance was initially identified through experimental genetic crosses between sensitive and resistant parasites^20,23,24^, and has also been suggested by genomic association studies^19,25–28^. Amambua-Ngwa et al. recently functionally characterised the role of PfAAT1 polymorphisms in resistance, demonstrating that the polymorphism S258L, present in *P. falciparum* isolates throughout Southeast Asia and Sub-Saharan Africa, increases chloroquine resistance at the expense of parasite fitness. They further demonstrate that F313S, which occurs alongside S258L in SEA, moderately reduces the S258L-mediated chloroquine resistance phenotype whilst partially restoring fitness. They propose F313S may help maintain chloroquine resistance associated genotypes (e.g. PfCRT), after the withdrawal of drug selection^19^. Potential co-operation between PfCRT and PfAAT1 is supported through further genomic studies^19,29^ and genetic crosses^19^.

Recent whole genome sequencing studies of Colombian *P. falciparum* isolates revealed the presence of additional novel polymorphisms in PfAAT1^25^. In light of the *de novo* emergence of chloroquine resistance in this region^7^ and the presence of unique PfCRT haplotypes in South America that are not found in Southeast Asia and Sub-Saharan Africa ^12^, we hypothesised that these uncharacterised polymorphisms could also modulate chloroquine sensitivity and/or fitness. We used structural modelling to explore the potential impact of these South American-focused polymorphisms on PfAAT1 structure, and then CRISPR/Cas9-mediated genome editing in lab-adapted *P. falciparum* strains of different geographic origins to functionally explore their impact on both chloroquine sensitivity and parasite growth rates. We suggest that a combination of PfAAT1 SNPs mediate complex fitness and sensitivity trade-offs, and identify a novel PfAAT1 SNP that is concentrated in South America and directly impacts chloroquine resistance profiles, suggesting it should be included in global molecular monitoring for drug resistance alleles.

## Results

### Frequency of AAT1 polymorphisms in South American isolates

Chloroquine resistance emerged historically completely independently in South America, where it has been maintained for multiple decades^25,30,31,33^, but the role of PfAAT1 variants in this emergence has not been extensively explored. We therefore focussed specifically on PfAAT1 sequences from South American *Plasmodium falciparum* isolates. Searching publicly available global whole genome sequencing data from MalariaGEN^14,34^ identified four SNPs in PfAAT1 that were found in South America but were not present in the reference African origin isolate 3D7; Table 1 shows their frequencies in both South American and global sample sets. The S258L polymorphism, previously characterised elsewhere^19^, is found in 45.9% of South American isolates, as well as in 64.1% of isolates globally. By contrast F313S, which has been reported to balance the fitness effects of S258L sensitivity^19^ was not found in any South American isolates, but was common in the broader global dataset.

**Table 1.** PfAAT1 polymorphisms and haplotype counts in different geographic regions. ^a^ Refers to *in vitro* adapted isolates (n=20) collected in Colombia which were genotyped in this study^14,32,33^. ^b^ Refers to isolates (n= 234) collected from Ecuador and Colombia and previously subject to whole genome sequencing^25^. ^c^ Data on haplotype prevalence obtained from MalariaGEN Pf8 online database for both the South America WHO Region (n=283, which includes many of the 234 Colombia/Ecuador genomes), and globally (n= 33,325)^14,34^. Percentages refer to the proportion of isolates within that sample set that contained a given polymorphism. Specific polymorphisms co-occur, and this explains why the total count under Global is greater than 100%. Conversely, the Ecuador-Colombia dataset contains some isolates which failed to successfully sequence at all loci, leading to a percentage total less than 100%.

|  | Colombia<sup>a</sup> | Ecuador-Colombia<sup>b</sup> | South America<sup>c</sup> | Global<sup>c</sup> |
| --- | --- | --- | --- | --- |
| V231D | 0 | 2.1% | 51.6% | 0.4% |
| I248T | 30% | 2.1% | 2.5% | 0.1% |
| S258L | 70% | 80.3% | 45.9% | 64.1% |
| P446A | 0 | 0 | 12.3% | 0.1% |
| F313S | 0 | 0 | 0 | 35.9% |

**Table 2:**
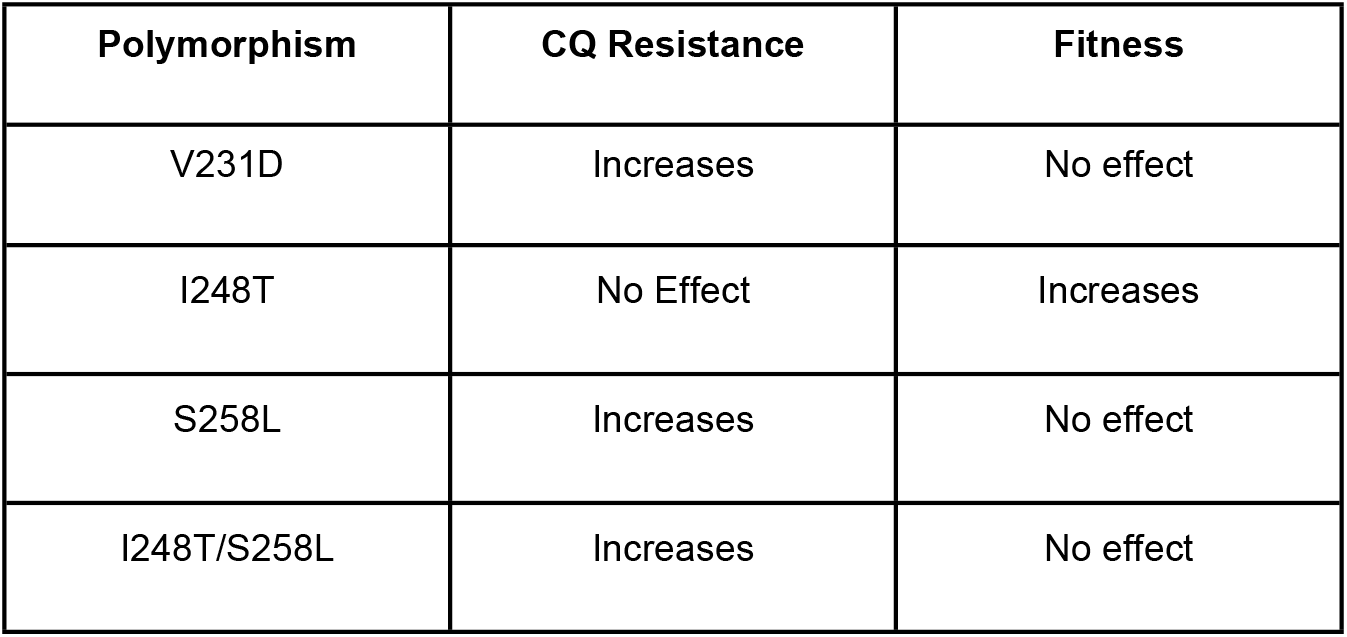
Summary table of 7G8 Polymorphisms and their experimentally determined resistance and fitness phenotypes.

| Polymorphism | CQ Resistance | Fitness |
| --- | --- | --- |
| V231D | Increases | No effect |
| I248T | No Effect | Increases |
| S258L | Increases | No effect |
| I248T/S258L | Increases | No effect |

The three other PfAAT1 SNPs were all concentrated in South America, and none have been characterised functionally. I248T, which is present on the same predicted transmembrane helix as S258L, is much more restricted and rare. As Table 1 shows, I248T is found in 30% of isolates from a panel of 20 culture-adapted Colombian isolates^32^, but only 2.5% of South American isolates and 0.1% of global isolates^34^, suggesting that parasites containing I248T are concentrated in Colombia. Another polymorphism, V231D, is present in 51.6% of isolates in South America as a whole, but only 0.4% of global samples. Interestingly, V231D is found in 7G8, a chloroquine-resistant line^41^ collected from Brazil in 1980^42^ and widely used as a reference culture-adapted strain for South American *P. falciparum*. Finally, P446A was found in 15 isolates exclusively in combination with V231D, and uniquely in South America. While a complete analysis of the correlation, or lack thereof, between PfAAT1 and PfCRT haplotypes is beyond the scope of this study, it is important to note that the PfCRT haplotypes common in South America, SVMNT (largely found in Brazil; also present in Oceania) and CVMET (common in Colombia) are not common in Africa or elsewhere, where either CVIET or wildtype haplotypes dominate.

### Structure predictions of PfAAT1 and potential impact of variants

To assess the potential structural and functional implications of PfAAT1 variants, we used AlphaFold to predict the overall structure for the protein (Figure 1a). The model shows that PfAAT1 possesses a highly disordered N-terminal region encompassing residues 1-169 followed by a compact multi-helical domain. The pLDDT scores for this model (Figure 1b) indicate that the prediction for the helical integral membrane region of PfAAT1 encompassing residues 170-606 is highly confident except for a short region that extends out of the membrane (residues 475-516). Using this model of PfAAT1 we mapped the position of the polymorphisms present in South American isolates, demonstrating they are all localised within the well-ordered helical bundle (Figure 1c). Visual inspection of the individual sidechain variants suggests that I248 helps stabilise the position of the loop and helix containing V231 (Figure 1d). Therefore, mutation of I248 to T, replacing a hydrophobic sidechain with a polar one, would disrupt local interactions potentially destabilising the structure and altering the conformation of the region encompassing V231. S258 forms inter-helical contacts within the core of the fold, some distance from the region containing I248T and V231D. Mutation of S258 to L, replacing a polar sidechain with a hydrophobic one, would be predicted to destabilise the fold of this domain (Figure 1e). P446 is also positioned within the core of the fold, within a helix. Mutation of P446 to A would likely impact the overall fold, due to the change from a rigid, cyclic sidechain to a more flexible backbone conformation (Figure 1f).

**Figure 1:**
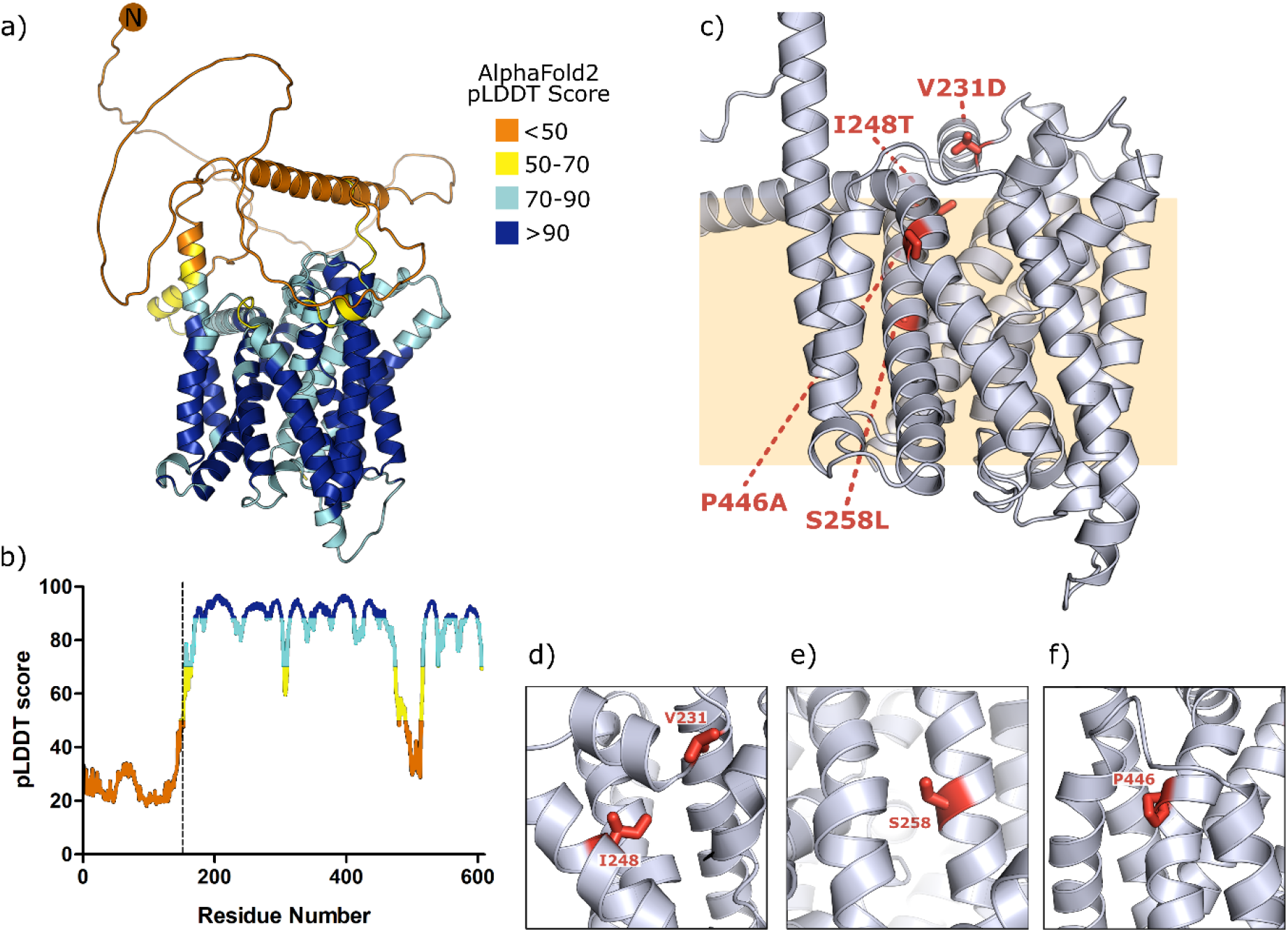
Structural predictions of PfAAT1 and polymorphisms. a) AlphaFold2 model of full-length PfAAT1 coloured by predicted local distance test (pLDDT) score. b) Plot of pLDDT score against residue number, with a dotted line delineating the N-terminal disordered region from the rest of the protein. The AlphaFold2 prediction for the structure of the N-terminus is of low confidence pLDDT<50. c) The positions of PfAAT1 polymorphisms (red) mapped onto the integral membrane portion of the predicted structure. Zoomed in images of the local interactions are illustrated for (d) I248 and V231, (e) S258, and(f) P446.

Detailed analysis of the predicted PfAAT1 structure identified a partially open channel accessible from one face of the membrane domain (Figure 2a). Structural homology searching using DALI^43,44^ identified a highly reliable structural homologue in humans, SLC38A9 (Z-score = 40.1, root mean square deviation RMSD = 1.25 Å, Figure 2b). SLC38A9 is a lysosomal amino acid transporter that senses luminal arginine levels, consistent with the known digestive food vacuole localisation of AAT1 which has similarities to a mammalian lysosome^45^. Furthermore, SLC38A9 possesses a partially open channel similar to that of AAT1 (Figure 2c) and a disordered N-terminus that has been suggested to form a plug that inserts into the pore at low arginine concentration to regulate transport^46^. This structural homology allows us to propose the orientation of AAT1 in the vacuolar membrane and that the disordered N-terminal region of AAT1 may sit within the cytosol and regulate transport of small molecules, potentially including amino acids, across the vacuolar membrane. Interestingly, V231 sits at the entrance to this opening suggesting that mutation to D may influence the transport of molecules such as chloroquine into or out of the vacuole. P446 is positioned within a helix close to the pore and could be important for the conformational changes involved in N-terminal plug activity. The mutation of P446 to A may therefore also impact transport across the vacuolar membrane.

**Figure 2.**
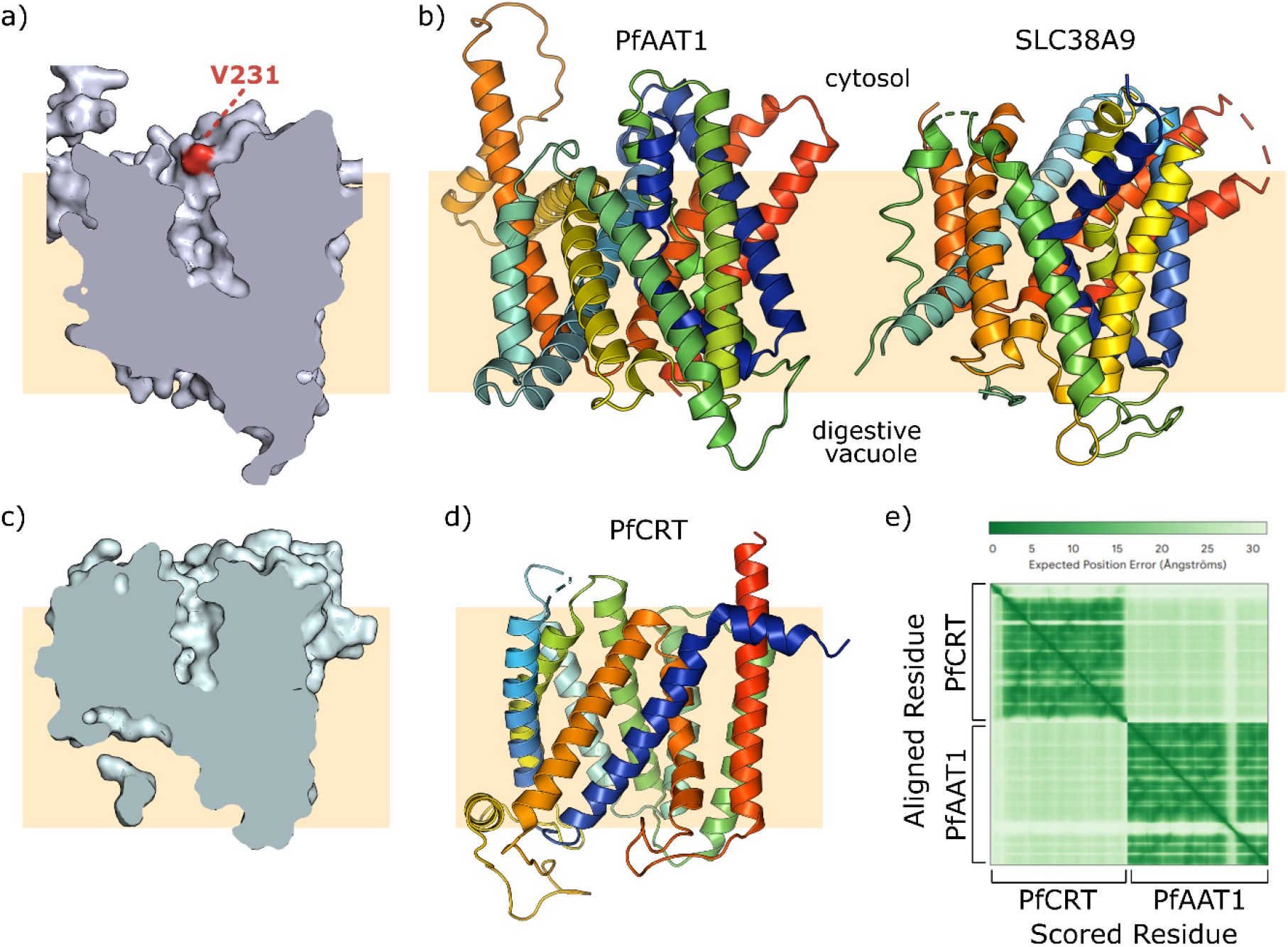
Structural homology of PfAAT1. a) Surface representation (grey) of PfAAT1 AF2 model and cut-through to demonstrate a partially open channel with V231 lying at the opening (red). b) Structural homology searching identified the human lysosomal arginine transporter SLC38A9. The PfAAT1 model and the SLC38A9 experimental structure (PDB ID: 7KGV) are shown as ribbons coloured from N-terminus (blue) to C-terminus (red) and illustrating their predicted membrane orientation. c) The structural similarity between these proteins extends to an equivalent partially open channel in SLC38A9. Surface representation and cut through of SLC38A9 in the same alignment as PfAAT1 in panel (a). d) The experimental structure of PfCRT^47^ (PDB ID: 6UKJ) does not possess significant structural similarity to the predicted structure of PfAAT1. e) The Predicted Aligned Error (PAE) plot of AlphaFold Multimer predictions of PfCRT-PfAAT1 complex formation suggest these proteins do not interact.

Given that PfAAT1 and PfCRT are both located in the digestive vacuole and are both genetically linked to chloroquine resistance, we also explored any structural homology between them and whether these two transporters might physically interact with each other. Although PfCRT also possesses a helical integral membrane domain, the overall structure is significantly different from PfAAT1 (Figure 2d) with a RMSD of 27.2 Å over 370 paired Cα atoms. Predictions of complex formation using Alphafold Multimer yielded proposed complexes that were extremely low confidence according to predicted alignment error statistics (Figure 2e). This suggests that PfCRT and PfAAT1 probably do not interact directly within the vacuolar membrane. Predictions using the sequence variants identified above had no impact on the PAE interacation scores. There is therefore no computational evidence that PfCRT and PfAAT1 interact physically, and their combined effects on chloroquine resistance are likely related to their related functions as vacuolar transporters.

### Generating transgenic PfAAT1 Lines

To explore potential roles of the South America-enriched PfAAT1 SNPs I248T and V231D on chloroquine sensitivity we used CRISPR/Cas9 genome engineering to modify these residues on a range of *P. falciparum* lab-adapted isolates. We included S258L in these studies as it sits in the same transmembrane region as I248T, but did not study P446A as it is always linked to V231D, and V231D is much more common (see Table 1). For the remainder of this work for simplicity we refer to these three SNPs as a single haplotype, with VIS representing the reference 3D7 haplotype, which is also present in the South-East Asian chloroquine resistant strain Dd2. The 3D7 haplotype of VIS is referred to as the reference haplotype throughout, with any differences from this haplotype indicated by bold residues. VI**L** representing the haplotype that has recently been functionally associated with resistance in African isolates^19^, and V**T**S (found in *in vitro* adapted isolates from Colombia) and **D**IS (found in 7G8), as the new haplotypes described in this study. Neither I248T nor V231D have ever been formally investigated for a role in mediating chloroquine resistance.

Isolates were selected to reflect a variety of PfCRT haplotypes as well as different geographical origins: Brazilian line 7G8, Colombian field isolate ColTu02.01^32^, Sub-Saharan African line 3D7^48^, and South-East Asian line Dd2^49^. The editing approach is shown diagrammatically in Figure 3a, and the successfully generated transgenic lines are summarised in Figure 3b. The chloroquine sensitivities of 3D7, 7G8 and Dd2 have been characterised extensively previously, most recently in work where we used them as comparators^32^. We obtained an IC50 for wildtype 7G8 of 70.2 nM, ColTu02.01 of 62.4 nM, Dd2 of 101.0 nM, whereas the IC50 value of wildtype 3D7 was over 10-fold more sensitive at 6.4 nM, reflecting its origin before chloroquine resistance emerged.

**Figure 3:**
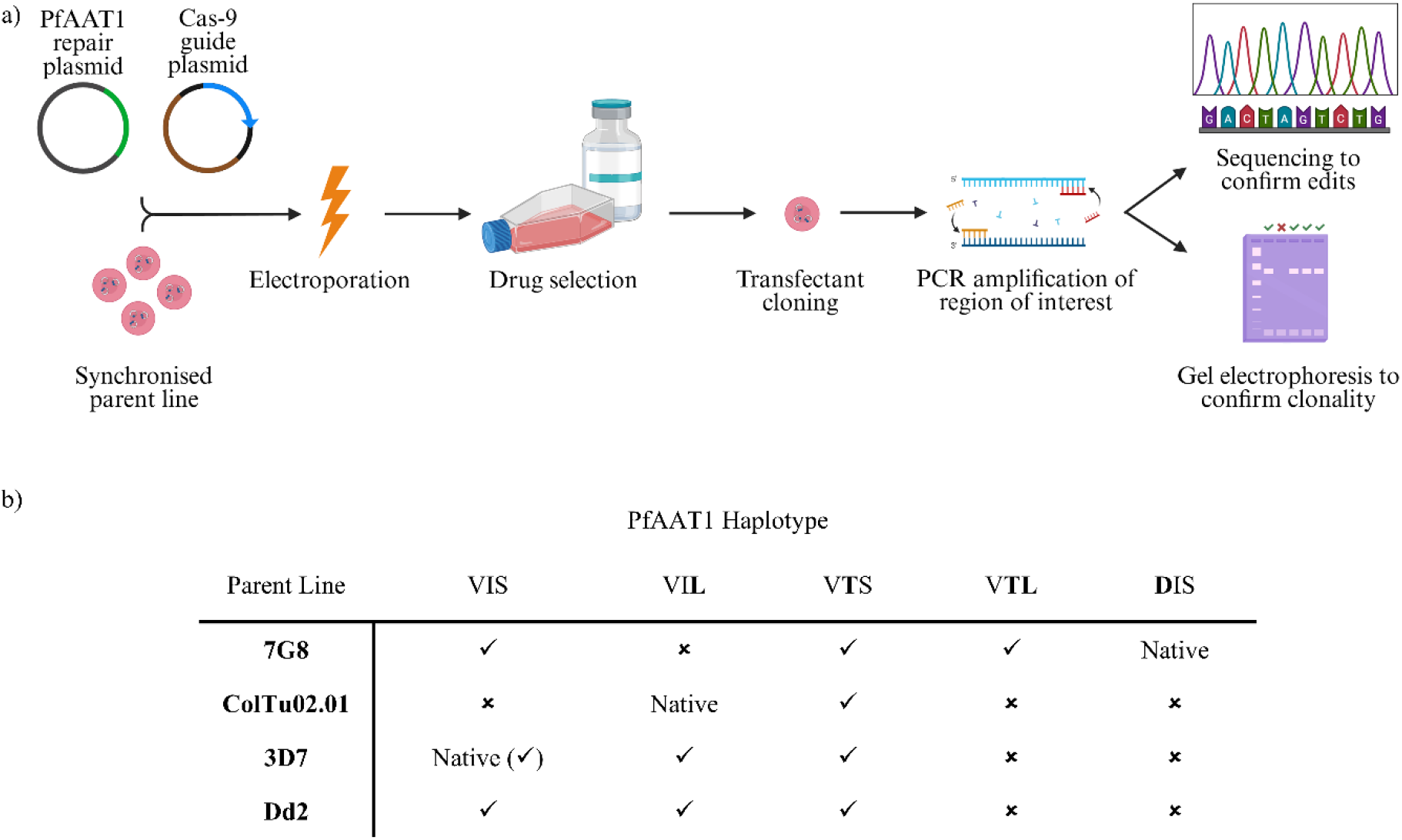
Generation of PfAAT1 transgenic *Plasmodium falciparum* lines. a) Schematic of generation of PfAAT1 transgenic lines, and confirmation using Sanger sequencing and gel electrophoresis. Created using BioRender.com b) Table detailing the specific combination of transgenic parasites created, listed by parent line on the first column, and PfAAT1 haplotypes on the first row. Native denotes the haplotype naturally present in that genetic background, while ticks denote successful generation of transgenic line, and crosses denote unsuccessful generation, often despite repeated attempts. 3D7 was transfected with a recodonised native sequence to serve as a transfection control, indicated by (tick). The native Dd2 haplotype is VIS**(F)**, containing the additional F313S polymorphism, but this sequence was not recreated by recodonisation; neither were the native 7G8 and ColTu02.01 sequences.

Some haplotypes were not generated successfully despite frequent independent transfection effects – for example we were unable to generate VI**L** on the 7G8 line (n=11 transfection attempts), despite the successful generation of VI**L**-expressing transgenic lines in both 3D7 and Dd2 backgrounds. This could potentially indicate a biological incompatibility of this haplotype with this specific genomic background, but given the high level of variability in *P. falciparum* transfections, technical limitations are the more likely explanation. We also noted that despite the fact that no *P. falciparum* isolates from South America have been identified that carry both I248T and S258L mutations, we were able to successfully generate this combination in 7G8 and it is observed naturally in Oceania-New Guinea.^34^

### Impact of altering PfAAT1 haplotypes on chloroquine resistance

The successfully generated transgenic lines were then assayed for CQ resistance using SYBR Green to quantitate parasitemia after *in vitro* growth in the presence of increasing concentrations of CQ. Editing 3D7 with a recodonised region that recreated the native 3D7 PfAAT1 VIS haplotype did not result in a significant difference in IC50 (Figure 4a; one-way ANOVA with Dunnet’s multiple comparison, p >0.05) confirming that neither the recodonisation nor the transfection process resulted in changes in drug resistance. Adding in either of the I248T or S258L mutations (resulting in V**T**S and VI**L** haplotypes respectively) had no significant impact on chloroquine responses (Figure 4a), indicating that alterations in PfAAT1 alone, in the absence of chloroquine resistance variants in PfCRT, is not sufficient to generate chloroquine resistance.

**Figure 4:**
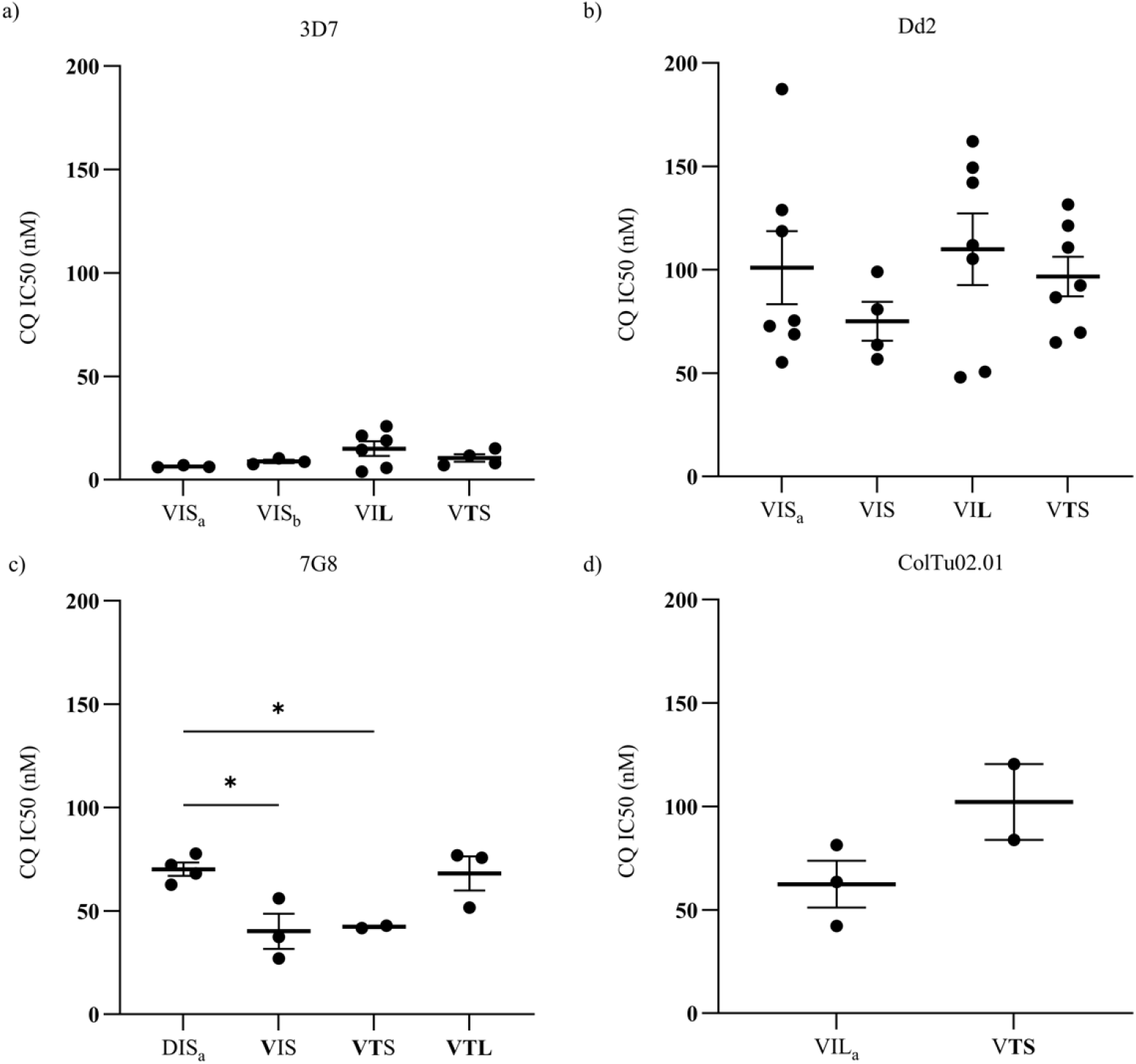
Chloroquine sensitivities of PfAAT1 transgenic lines. Charts displaying biological replicates of chloroquine sensitivity assays of CRISPR-Cas9 edited lines expressing transgenic PfAAT1 sequences. The IC50 value is expressed on the y axis, PfAAT1 haplotype on the x-axis, and parent line displayed above, corresponding to a) 3D7, b) Dd2, c) 7G8, and d) ColTu02.01. Sensitivity data for the unedited wildtype is shown by the haplotype denoted with subscript a. Recodonised wildtype is denoted by subscript b. Mutant polymorphisms, relative to wildtype, are highlighted in each haplotype in bold. Mean and standard error of the mean shown. Significance calculated for a, b & c using one-way ANOVA, assuming gaussian distribution and same standard deviation, with corrections for multiple comparisons relative to wildtype parasite. Significance calculated for d using Unpaired T-test. Statistical analysis and chart plotted using GraphPad, Prism.

In Dd2, there is an additional polymorphism, F313S, which has been predicted to be associated with fitness and resistance in different lines from Southeast Asia^19^. However, we observed no significant difference between the chloroquine sensitivity of the Dd2 parent line, VIS(S), and a Dd2 strain in which the F313S mutation has been edited out, VIS(**F**) (Figure 4b). These data therefore do not support a significant role of the F313S polymorphism, which had been proposed previously, at least in this background^19^. Editing Dd2 PfAAT1 SNPs also had no impact on chloroquine sensitivity in the Dd2 background, as again converting VIS to V**T**S or VI**L** resulted in no change in drug response (Figure 4b). This may reflect a differential role of transporter proteins in mediating the resistance phenotype, as proposed previously,^18^ or some association between the phenotypic impact of variants and PfCRT haplotypes. Notably, both 7G8 and ColTu02.01 contain different PfCRT haplotypes (SVMNT and CVMET, respectively,) to Dd2 (CVIET).

By contrast, in the 7G8 South American background modifying PfAAT1 SNPs did affect chloroquine sensitivity (Figure 4c). Reverting the V231D mutation, i.e. converting the D normally found at residue 231 to V caused a significant decrease in IC50 value relative to the wildtype line (one-way ANOVA with Dunnet’s multiple comparison, p = 0.019). Editing out V231D, i.e. converting **D**IS to VIS, therefore decreases chloroquine resistance in 7G8, supporting a role for this variant in mediating chloroquine resistance. This reduction in chloroquine resistance was maintained when the I248T was added to create the V**T**S haplotype (Figure 4c; one-way ANOVA with Dunnet’s multiple comparison to the native DISF haplotype, p = 0.048), which suggests that I248T has no additive role in this background. By contrast, in the absence of V231D but the presence of both S258L and I248T, creating the V**TL** haplotype, chloroquine resistance was restored to background 7G8 levels (Figure 4c; no significant change in IC50 value in the V**TL** transgenic line compared to parent line, one-way ANOVA with Dunnet’s multiple comparison, p >0.05). In 7G8 therefore, both the V231D and S258L polymorphisms are able to generate a chloroquine resistance phenotype. This is the first time that PfAAT1 residue V231 has been functionally implicated in chloroquine resistance. Given the common frequency of this variant in South America, and relatively much lower prevalence in other regions, the emergence of V231D could be hypothesised to have played some part in the origin of CQ resistance in South America^34^.

ColTu02.01 is a chloroquine resistant Colombian field isolate collected from the Pacific Coast department, Nariño, in 2002, and contains the PfAAT1 haplotype VI**L**. After successful long-term adaptation of this isolate to *in vitro* culture^32^, we successfully edited the PfAAT1 locus to convert the native VI**L** haplotype to V**T**S. There was no significant difference in chloroquine sensitivity (unpaired T-test, p > 0.05) between the parent and transgenic line (Figure 4d). V**T**S can therefore substitute for VI**L** in this background, suggesting for the first time that I248T, which seems largely restricted to Colombia, may also play a role in CQ resistance.

All transgenic lines were also assayed for mefloquine sensitivity, but there was no significant change in resistance between parent lines and any of the respective transgenic lines (supplementary figure 1.)

### Modification of PfAAT1 residue 248 in the 7G8 background has a fitness effect

Drug resistance-associated polymorphisms frequently also influence fitness. We therefore used non-competitive growth assays to assess whether any PfAAT1 changes had an effect on fitness (Figure 5a), selecting the transgenic lines created in 7G8 as this is the background in which we had been able to generate the widest range of haplotypes and also had the clearest differences in sensitivity phenotype between haplotypes. Alteration of amino acid 231 alone had no impact on growth as measured by Parasite Multiplication Rate (Figure 5b; PMR; comparison of VIS transgenic line and 7G8 parent line with **D**IS haplotype using one-way ANOVA with Dunnet’s multiple comparison, p >0.05.) However, both the V**T**S and V**TL** transgenic lines had an increased Parasite Multiplication Rate (PMR) when compared to 7G8 parent line (DIS), with the V**T**S line reaching significance (one-way ANOVA with Dunnet’s multiple comparison, p = 0.012). This proposes a model wherein I248T increases the fitness of parasites within a 7G8 genomic background. S258L has been proposed to incur fitness costs^19^, and this may explain the absence of a significant increase in PMR of triple mutant **VTL**, relative to parental line DIS. I248T may therefore not be capable of off-setting the entire fitness cost of S258L.

**Figure 5:**
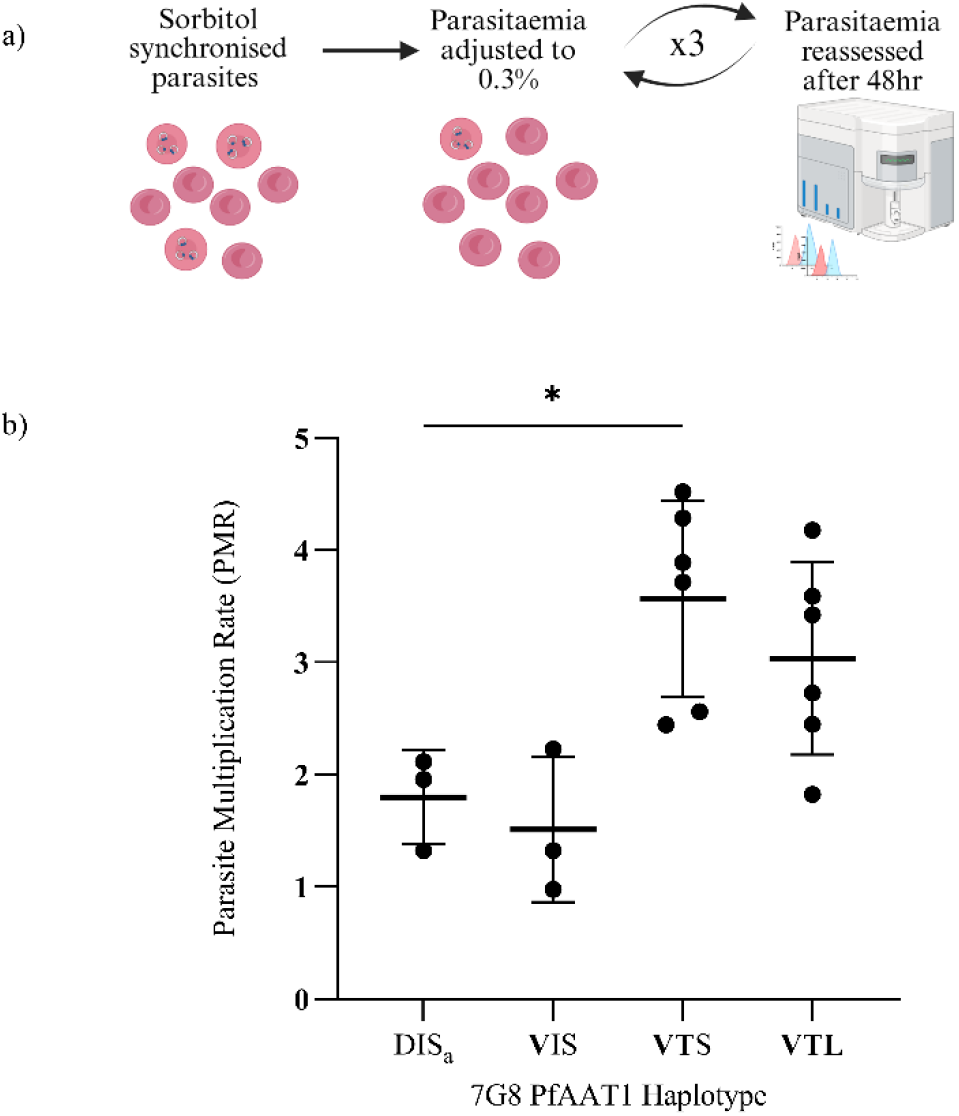
Growth Assay of 7G8 Transgenic Lines. a) Diagram of growth assay set-up. Sorbitol synchronised transgenic lines were grown in triplicate, and their parasitaemia measured before and after each growth cycle using a flow cytometer, to calculate Parasite Multiplication Rate (PMR). This was repeated for 3 growth cycles. Growth data for the unedited wildtype is shown by the haplotype denoted with subscript a. Figure made with BioRender.com b) Chart of the PMR of the 7G8 transgenic lines, and the 7G8 unedited line. Significance calculated with one-way ANOVA, assuming normal distribution and equal standard deviation, and with multiple comparisons relative to wildtype 7G8.

## Discussion

While the first licensed *Plasmodium falciparum* vaccines have the potential to make a significant impact on malaria globally^50^, their rollout is still in an early stage and antimalarial drugs, rapid diagnostic tests and insecticides remain the mainstay of global control efforts^51^. Antimalarials have without question saved millions of lives but their history is one of repeated advance and retreat, with the development and release of new drugs being repeatedly counteracted by the evolution and spread of parasite drug resistance^52^. Understanding the functional causes of resistance is therefore of critical importance. This is particularly true as resistance first emerges, where causative associations can help inform genetic surveillance as well as the more precise targeting of malaria control efforts. However, understanding the molecular causes of resistance is important even after resistance is widespread, as the parasite fitness costs that often accompany resistance can mean that resistance variants become selected against and decrease in frequency when the drug is no longer in common usage. Such is the case for chloroquine resistance, where resistance alleles seem to have largely disappeared from certain regions now that CQ is no longer widely used, raising the potential that such drugs could be re-used as part of a coordinated and data-informed malaria control strategy^53,54^.

Chloroquine (CQ) was the first truly globally used antimalarial drug and it formed the basis of widespread malaria eradication efforts in the 1950s and 1960s^55^. This global selection pressure resulted in the emergence of CQ resistance in the 1960s, with resistance evolving independently in South East Asia, the South Pacific and South America before spreading to Africa^56^. Pioneering cutting-edge molecular studies, carried out long before genetical manipulation of the *P. falciparum* genome became routine, identified variants in the food vacuole transporter PfCRT as the primary cause of CQ resistance^12,49,57,58^, with the K76T variant associated with CQ resistance on a wide range of different genomic backgrounds and haplotypes^12^. However, it is now becoming increasingly apparent that CQ resistance is multigenic, and a second food vacuole transporter PfAAT1 plays an additional modulatory role. Two PfAAT1 SNPs, S258L and F313S have been functionally shown to modulate CQ resistance and parasite fitness, but these studies have only been carried out in the context of African and Southeast Asian parasites^19^. This manuscript highlights three additional PfAAT1 SNPs (V231D, I248T and P446A) which form the major PfAAT1 polymorphisms in South America, following S258L. Of these three, V231D is the most common, being present in nearly 50% of isolates in South America. P446A is only found in combination with V231D and only in a minority of isolates, while I248T seems largely confined to Colombia.

Mapping these variants onto a highly robust AlphaFold model of PfAAT1 illustrates how the different residues may influence the overall fold of the protein and provides information about their relative positions. Residues 231 and 248 are close in the primary amino acid sequence, and analysis of the structure demonstrates how the mutation I248T may alter the conformation of the loop containing V231. S258 is also quite close in the primary sequence, however it is structurally more distant from residues 231/248. This emphasises the limitations of functional predictions based on primary amino acid sequence or two-dimensional topologies, and the benefit of confident structural models in interpreting variants identified in genome-wide association studies.

We also identify a strong structural homology between PfAAT1 and the arginine transporter, SLC38A9, which allows prediction of the orientation of PfAAT1 within the food vacuole membrane and reveals further functional insight. V231 is present on the proposed cytoplasmic face of PfAAT1 at the gateway of a partially open channel that may be plugged by a flexible N-terminal domain. Therefore, it is possible that variation at this position could affect transport through the pore, and therefore potentially negatively influence uptake of chloroquine and/or other molecules into the food vacuole. By contrast I248T and S258L sit within the same transmembrane helix that forms the channel within the food vacuole membrane, but given the role of I248 in stabilisation of the helix and loop on which V321 resides, mutation of I248T may also affect the capacity of PfAAT1 for transport.

While this manuscript focuses on South American variants of PfAAT1, the insights that structural modelling generated led us to map all 14 polymorphisms observed to date in the Pf8 global genome sequencing dataset^14^ (Figure 6a). The second most common variant globally, after S258L, is Q454E and is almost exclusively seen in isolates from Southeast Asia (AS-SE-E). Like V231D, this residue sits over the open pore of PfAAT1 and introduces an acidic charge; both D and E being negatively charged sidechains (Figure 6b). Another variant, G308E, with a broader global distribution, also lies near the open pore of PfAAT1 and again introduces a negative charge. The shared molecular localisation and negative charge of these three variants suggest they may all contribute in a similar way to altering the transport function of PfAAT1. The function of the PfAAT1 structural homologue SLC38A9 in humans is to transport arginine residues, which are positively charged, out of the lysosomal compartment and into the cytosol^59^. It is therefore tempting to speculate that the variants in AAT1 that enhance the negative charge at the cytosolic face of this transporter may facilitate more effective efflux of small positively charged molecules such as arginine and drugs, including CQ. In further support of an electrostatic driver playing a role in selection for PfAAT1 variants, K541N, which is seen across Asian isolates, sits on the opposite, vacuolar side of PfAAT1and removes a positive charge from this face, further enhancing the potential drive to increase export of positively charged small molecules from the vacuole (Figure 6c).

**Figure 6:**
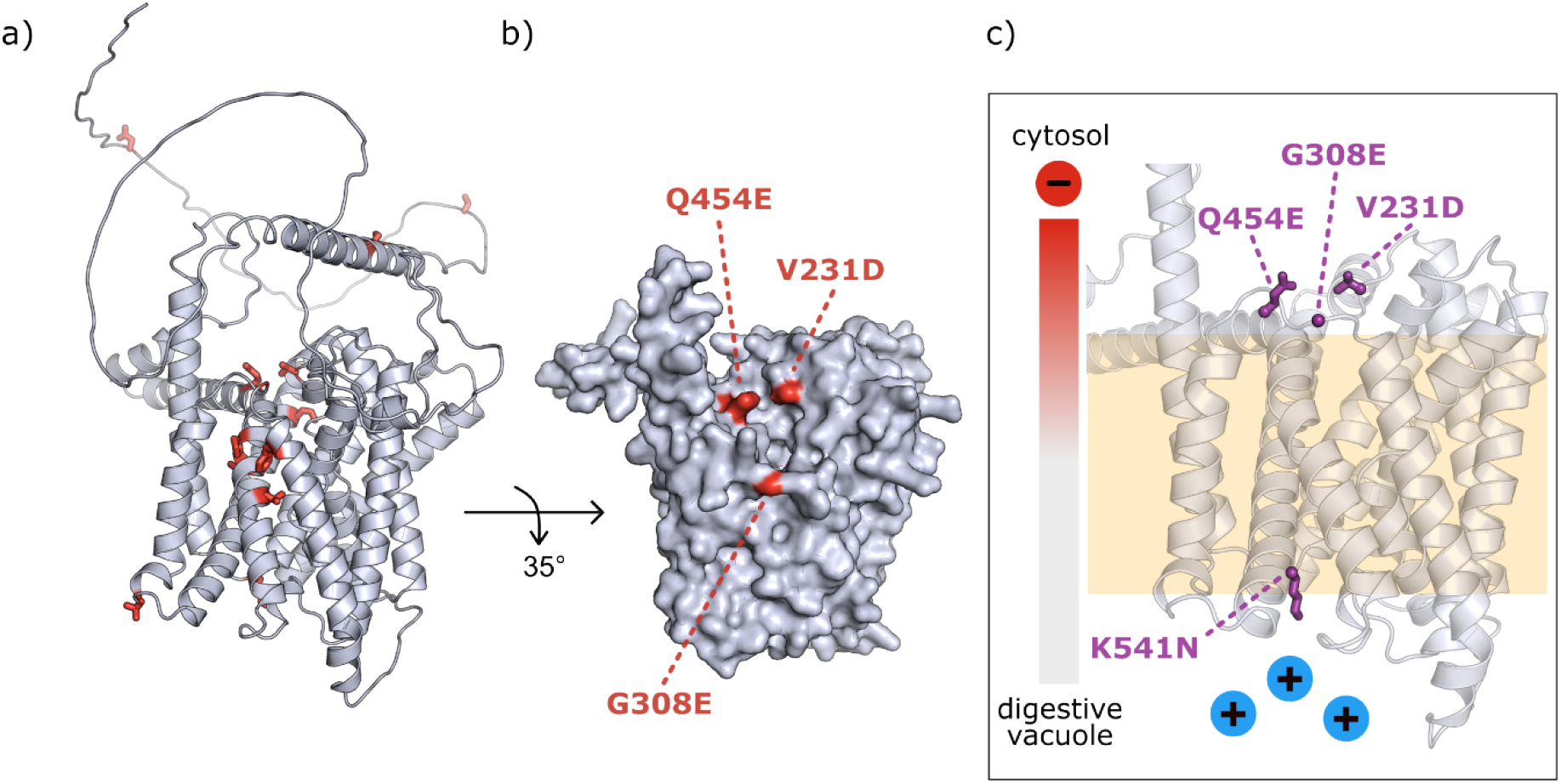
Global PfAAT1 polymorphisms. a) The positions of all current polymorphisms identified in MalariaGEN Pf8^14^ (D17N, S43I, I61L, V231D, I248T, T282N, G308E, F313S, T440A, V445I, P446A, Q454E, K541N; red) were mapped onto the AlphaFold2 prediction of full-length PfAAT1. Several haplotypes contain one or two of these mutations. b) Surface representation (grey) of the integral membrane portion of the predicted structure of PfAAT1, rotated by 35° relative to panel (a). Amino acids V231, G308, and Q454 are highlighted (red) illustrating their proximity to the open pore on the cytosolic face. c) Schematic diagram showing the electrostatic gradient (left) generated by the SNPs V231D, G308E, Q454E, and K541N (purple). These mutations may support the efflux of positively charged small molecules (blue) such as drugs or amino acids from the food vacuole towards the enhanced negative charge (red) at the cytosolic face of PfAAT1.

Most of the other globally observed polymorphisms occur within the transmembrane portion of PfAAT1 are potentially destabilising, involving changes in the polarity of amino acids that make inter-helical contacts within the core fold. Three of the 14 SNPs (D17N, S43I, I61L) are within the disordered N-terminus of the predicted structure, which homology to SLC38A9 suggests may have a role as a regulatable plug for the pore. The precise functional impact of these SNPs on this putative plug are not possible to predict. Finally, why certain mutations occur together within haplotypes is difficult to fully explain from structural modelling, but may reflect either genetic-linkage effects or potential long-range compensatory effects.

CRISPR-Cas9 engineering confirmed the functional insights from structural models regarding South American variants. In the 7G8 background removing the V231D mutation decreases CQ resistance almost 2-fold, showing for the first time that PfAAT1 V231D is involved in chloroquine resistance. I248T alone was not sufficient to return CQ resistance to wild-type levels, but when combined with S258L in the **VTL** haplotype, 7G8 CQ resistance is restored. This supports the previously established role of S258L as a mediator of chloroquine resistance on a mutant PfCRT background. Interestingly I248T appeared to have a significant fitness effect in the 7G8 background. We therefore hypothesise that I248T has emerged in some South American parasites as a potentially balancing variant to counteract the fitness cost of mutant PfCRT.

Our data also emphasise the importance of geographic context. Modifying I248T or S258L had no impact on CQ resistance in the Southeast Asian Dd2 background, although resistance data was in general more noisy in this line. We were unable to generate V231D variants in Dd2, but as noted in the Results section this is very likely to reflect technical challenges and should not be interpreted as implying that this variant is not tolerated in a Dd2 background. However, the different phenotypic impacts of the variants in different *P. falciparum* lines, along with the geographical restriction of the variants themselves, all clearly indicate some genomic background-dependency of the effect of PfAAT1 variants. The most likely cause of this is PfCRT, as PfCRT haplotypes in South America in general, and Colombia in particular, are well-established to be different to those that emerged in Southeast Asia^14,34,60^. Similarly, no PfAAT1 mutations were sufficient to cause CQ resistance in the sensitive 3D7 background, emphasising again that PfAAT1 plays a modulatory role to PfCRT, and is unlikely to cause CQ resistance alone. More comprehensive studies of combinations of different PfAAT1 variants in different genomic backgrounds and different PfCRT haplotypes are clearly required to understand the full extent of the genetic interactions between PfAAT1 and PfCRT.

What are the implications of this for drug policy and genomic surveillance? For the former the implications are limited as CQ is no longer in widespread use, although they do suggest that population-level PfAAT1 variation should be considered alongside PfCRT variation if and when the reintroduction of CQ is ever considered at a local level. For surveillance however, whether through whole-genome sequencing (WGS) or targeted SNP amplicon panels, the implication is very clear – that V231D should join S258L as validated mediators of CQ resistance and be targeted for surveillance efforts, particularly in the context of South American isolates. In addition, we believe that this work emphasizes the importance of bringing together genetic, structural and functional data. WGS has revolutionised the study of antimalarial drug resistance, enabling the rapid identification of SNPs that increase in frequency as clinical resistance increases. Artemisinin resistance is a perfect example as the development of *P. falciparum* WGS coincided with the emergence and spread of artemisinin resistance, and a huge amount of effort has focused on tracking SNPs that associate with the frequency of resistance ^61–64^. However, it is always important to remember that these are associations alone, and further functional validation through direct genome engineering can be crucial in distinguishing driver from passenger mutations. *Plasmodium falciparum* genome engineering studies are still far from trivial however, and comprehensively exploring different combinations of variants and genomic backgrounds is extremely challenging. Here structural biology can be hugely useful, as our data shows that robust models can provide insight that primary sequence data alone cannot, and highlight specific residues or haplotypes to target. A more joined up approach, which brings together genomic surveillance, structural biology and experimental genetics, ideally based in malaria endemic countries, should be a priority for both researchers and funders.

## Materials and Methods

### Prevalence of Polymorphisms in PfAAT1

The prevalence of polymorphisms in Colombia was calculated using a panel of 20 historical and contemporary isolates. Historical isolates were collected in previous studies^30,31^ and have been extensively characterised^32,33^, and contemporary isolates, collected after 2006^32^, were sequenced using Sanger sequencing with GeneWiz, at PfAAT1 with primers ACACCTGGTGGTGTTAGATCTA and GATTTGGTTTGTAGTCCTATAGTTAATACAGT, amplified using PCR for 25 cycles at 49°C annealing temperature, and with an extension time of 30 seconds at 65°C. The historical isolates were additionally characterised in another study^25^ along with further isolates collected from Ecuador and Colombia. MalariaGEN datasets^14^, explored using PfHaplo-Atlas^34^, also included the historical samples characterised by these sources, but the application of different quality control metrics for variant calling means that only some isolates overlap across the different sources. Notably, some haplotypes contain >1 polymorphisms, e.g. I248T/S258L, and therefore these isolates were represented twice in the table, providing a separate count for I248T and S258L polymorphisms.

### Protein Structure Models

Sequences of PfAAT1 and PfCRT were taken from PlasmoDB (Gene ID: PF3D7_0629500 and PF3D7_0709000 respectively). Predicted structures of PfAAT1 were generated using AlphaFold2, AF3 and AF Multimer with all yielding the same overall fold^35,36^. Additional predictions using truncations that remove the unstructured N-terminal regions of PfAAT1 also yielded the same fold for the integral membrane portion. AlphaFold data are available as pdb files in the Supplementary Information. Models were analysed using WinCOOT^37^ and PyMol (The PyMol Molecular Graphics System, Version, 1.2r3pre, Schrödinger, LLC). Figures were generated using PyMol and ChimeraX^38^.

### CRISPR-Cas9 Genetic Editing

CRISPR-CAS9 guide, and donor template plasmids, were designed using Benchling software. The guides used for insertion to PfAAT1 were either GAAATTAAATACATAAAAGA^19^ or ACATTATTTGTGAATAAGGT, the latter designed using Benchling guide prediction software. The plasmids were amplified by transformation into *E. coli* XL10-Gold Ultracompetent Cells (Agilent) and successful integration was confirmed by diagnostic restriction digest, and Sanger sequencing across the insert region. Colonies expressing successfully edited plasmids were expanded, and the plasmid purified using mid-range plasmid purification kits (XtraMidi Plus, Nucleobond). Then 70 µg of donor template, and 30 µg of guide plasmid were precipitated, and resuspended in 30 µl Cytomix solution (120 mM KCl; 0.15 mM CaCl_2_; 10 mM KH_2_PO_4_; 25 mM HEPES; 2 mM EDTA; 5 mM MgCl_2_ at pH7.)

*P. falciparum* cultures were sorbitol synchronized, and 96 hours later were electroporated at the ring stage with the Cytomix solution containing the plasmid using GenePulser Xcel (BioRad) at 310 V, infinite resistance and 950 µF resistance. The transfected cultures were selected using the DHODH inhibitor DSM1 at 1.5 µM (Sigma-Aldrich, 5.33304). The DSM1 resistance cassette was contained on the guide plasmid.

Positive transfectants were cloned using limiting dilution cloning, involving a 10-fold serial dilution to reach a final dilution of 0.3 parasites per well. These were left for 11 days, and the formation of single plaques in a well assessed using 4X lens using the brightfield on EVOS FL imaging system (Thermofisher)^39^. The clonality of the parasite was confirmed by amplification and gel electrophoresis of the edited and unedited PfAAT1 region. The identity of the transfectant was confirmed using Sanger sequencing of recodonised sequence using GeneWiz with the following primer pair-GAACAAACAAACCAAAAAGAAACTGGAAAGGACGGACCTT, and AATTGTCCAGTAATAAAACAGGCGTTCGGCTGCTGGCTA. The sequences were analysed using Benchling software. Where possible, 2 clones were used per transgenic line.

### *In vitro* Drug Sensitivity Assays

*In vitro* sensitivity assays were carried out using a protocol provided by Prof. Marcus Lee, as per previous work^32^. Briefly, these involved sorbitol synchronisation of parasite cultures, and in the following cycle, the synchronised cultures were adjusted to 0.5% parasitaemia and 1% haematocrit. The diluted culture was then added along a serial drug dilution ran in triplicate across a 96-well plate, of either mefloquine (Sigma-Aldrich, M2319) or chloroquine (Sigma-Aldrich, C6628). This set-up was repeated but with a dilution series of the vehicle solution into which the drug was diluted as a further control. There were further controls of infected and uninfected erythrocytes grown in the absence of drug. These plates were incubated in a gassed incubator at 37°C for 72 hours, following which they were frozen at -20°C for > 24 hours.

The frozen plates were thawed, and incubated at 37°C for 2-4 hours with lysis buffer (20 mM Tris-HCl, 5 mM EDTA, 0.1% w/v saponin and 1% v/v Triton X-100 in MiliQ water) and 10,000X SYBR Green dye. SYBR Green acts as a nucleic acid stain, and fluorescence therefore indicates parasite growth. The plates were then read on a plate reader (BMG CLARIOstar) with 485 nm excitation, and 535 nm emission filters.

The starting chloroquine and mefloquine concentrations differed between isolates, reflecting both different predicted IC50s and data from subsequent optimisations, and ranged across 3D7 (1000-3000 nM (CQ) and 500 nM (MQ)); 7G8 (3000 nM (CQ) and 125-500 nM (MQ)); Dd2 (3000 nM (CQ) and 500 nM (MQ)) and ColTu02.01 (3000 nM (CQ) and 250-500nM (MQ)).

The results were exported to GraphPad Prism for non-linear regression. Two 96-well plates were set up for each day, and the mean of their two IC50 values formed a single biological replicate. If one plate failed, then a single plate was used as a biological replicate. The biological replicates for each transfectant line were first processed using outlier analysis, using ROUT method (Q>10%), and the subsequent data sets compared using one-way ANOVA test, assuming standard deviation and normal distribution, with post-hoc comparisons of each isolate’s mean against the parent line using Dunnet’s Multiple Comparisons Test.

### Non-Competitive Growth Assays

The non-competitive growth assay was adapted from a protocol provided by Emma Kals^40^. Parasite lines were thawed and cultured for 2 weeks. The cultures were sorbitol synchronised and then diluted to 0.3% parasitaemia in 2 mL. Each line was run in triplicate, and starting parasitaemia confirmed using Attune NxT acoustic focusing cytometer (Invitrogen). After 48 hours, the parasitaemia was rechecked, and the cultures readjusted to 0.3%. This was repeated for 3 cycles.

The parasitaemia was assessed by exporting fcs files to FlowJo software (www.flowjo.com/flowjo/overview), and identical gating applied for each replicate in a given 48 hour period. The average fold increase across the three replicates was designated the parasite multiplication rate, and the means were compared using a one-way ANOVA test, assuming standard deviation and normal distribution, with post-hoc comparisons of each isolate’s mean against parent line, using Dunnet’s Multiple Comparison Test.

## Ethics statement

Parasites were grown in human blood and serum sourced from NHS Blood and Transfusion (NHSBT). Informed consent for use of blood materials in research was carried out by NHSBT, and all samples were anonymised before transfer. The use of human blood products for this research was reviewed and approved by NHS Research Ethics Committee (approval 20/EE/0100).

## Acknowledgements and Data Availability

We thank Angela Early (Broad Institute) for help with obtaining allele frequencies for PfAAT1 SNPs and Richard Pearson (MalariaGEN) for insightful discussions on AAT1 and CRT haplotypes. Thank-you to the parasitology laboratory staff at the Universidad del Valle for their help and advice in preparing reagents and solutions, with a specific thanks to Professor Maria del Pilar Crespo for advice regarding the *in vitro* culture procedures, and the supportive staff at the Hospital Universitario del Valle. We extend our thanks also to all members of the Rayner laboratory in the University of Cambridge for their advice and support in the laboratory. The authors gratefully acknowledge Centro Interanacional de Entrenamiento e Investigaciones Medicas (CIDEIM) for providing *P. falciparum* Colombian samples under the Material Transfer Agreement – MTA # 048 and 056.

This work was supported by a Wellcome Trust Investigator award to J.C.R [220266/Z/20/Z], and received financial support by internal calls CI 11308 and CI 11299 from the Universidad del Valle, Cali, Colombia, to D.F.E. EB is funded by the University of Cambridge School of Clinical Medicine Doctoral Training Programme in Medical Research. J. E. D. was funded by a Wellcome Trust Senior Research Fellowship (219447/Z/19/Z)

For the purpose of open access, the authors have applied a CC BY copyright license to any Author Accepted Manuscript Version arising from this submission. The funders played no role in the study design, data collection and analysis, decision to publish, or preparation of the manuscript. There are no financial, personal or profession conflicts of interest to declare.

---

**Supplementary Figure 1:**
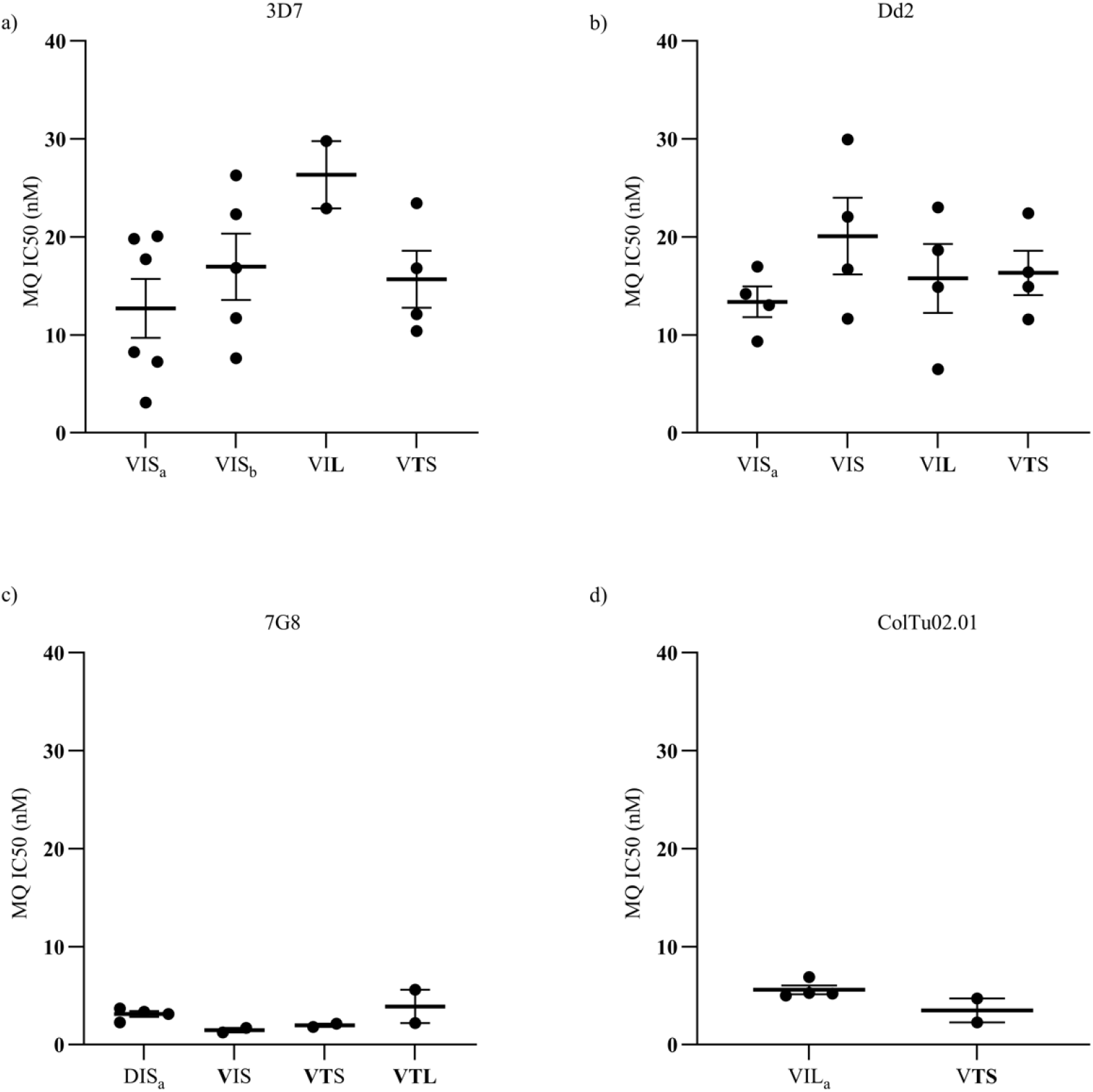
Chart of mefloquine sensitivities of PfAAT1 transgenic lines. Charts displaying biological replicates of mefloquine sensitivity assays of CRISPR-Cas9 edited lines expressing transgenic PfAAT1 sequences. IC50 value expressed in on the y axis, and PfAAT1 haplotype expressed on the x-axis, and parent line displayed in grey font behind the chart, corresponding to a) 3D7, b) Dd2, c) 7G8, and d) ColTu02.01. Sensitivity data for the unedited wildtype is shown by the haplotype denoted with subscript a. Recodonised wildtype is denoted by subscript b for 3D7 only. Mutant polymorphisms, relative to wildtype, are highlighted in each haplotype in bold. Mean and standard error of the mean shown. Significance calculated for a, b & c using one-way ANOVA, assuming gaussian distribution and same standard deviation, with corrections for multiple comparisons relative to wildtype parasite. Significance calculated for d using Unpaired T-test. Statistical analysis and chart plotted using GraphPad, Prism.

## Notes

### Competing Interest Statement

The authors have declared no competing interest.

